# Epinephrine Enhanced Fear Conditioning under Pentobarbital or Dexmedetomidine Anesthesia in Rats

**DOI:** 10.1101/2025.04.30.651393

**Authors:** K.-H. Chen, S.-T. Chao, H.-Y. Hsiao, S.-D. Chang, D.-Y. Chen, K.-C. Liang

## Abstract

This study examined effects of epinephrine injected in learning under anesthesia on awake memory of three fear conditioning tasks. Epinephrine given after training at various doses did not affect conditioned freezing acquired in an awake state or under 50 mg/kg pentobarbital; however, 0.1 mg/kg epinephrine caused saving in re-learning. For conditioned-fear potentiation of startle acquired in an awake state, epinephrine at 0.01 or 0.1 mg/kg enhanced 1-trial learning but at 1.0 mg/kg impaired 5-trial learning. For this task trained under pentobarbital, epinephrine enhanced 1-trial learning at 1.0 mg/kg and 5-trial learning at 0.1 or 1.0 mg/kg. Under infusion of dexmedetomidine (0.1 mg/kg/hr), learning of this task at a 0.63 mA foot shock yielded poor memory, which was improved by 0.1 mg/kg epinephrine; yet epinephrine had no effect on the better memory yielded by 1.25 mA foot shock. In an inhibitory avoidance task, rats in an awake state explored the apparatus and then received foot shocks under anesthesia of dexmedetomidine, injections of 0.1 mg/kg epinephrine before each shock training session enhanced memory. Deleting the awake exploration abolished the epinephrine effect. These results, taken together, suggest that epinephrine could enhance learning in three conditioning tasks under two anesthetics when the fear memory was assessed indirectly through its modification of an innate or learned behavioral tendency.

## Introduction

Significant events arousing emotion often leave indelible memory traces. This led to a notion that memory formation requires not only specific sensory inputs but also nonspecific arousal (Hebb, 1955; Konorski, 1967). Consistent with this proposal are findings that conditioned arousal was observed at an initial stage of conditioning and it played a critical role in developing the specific conditioned response (Pavlov, 1927; Lennartz and Weinberger, 1992). Thus, manipulation of neural and endocrine systems implicated in emotion or arousal functions during or shortly after learning was shown to facilitate acquisition/retention in many tasks, such as hypothalamic stimulation may accelerate eye-blink conditioning (Kim et al., 1983) and administration of some stress hormones including epinephrine could modulate memory consolidation (McGaugh, 1983).

The role of nonspecific arousal in memory formation bears upon an interesting question of whether memory could be acquired during sleep or anesthesia (Reasor and Poe, 2008). While memory processing during sleep is well documented (Inostroza and Born, 2013; Klinzing et al., 2019), learning under anesthesia remains a controversial issue. Some human studies reported that after resurrection anesthetized patients could remember events or materials presented during surgery when subjected to an implicit memory test (Merikle and Daneman, 1996). A study on rats about social transmission of food safety showed olfactory learning along with increase of unit activity in the olfactory bulb under anesthesia of isoflurane (Nicol et al., 2014).

Research also noted a role of nonspecific arousal in learning under anesthesia. In a priming task, humans performed better for words with negative emotion tones than neutral ones presented during anesthesia (Gidron et al., 2002). An early study trained anesthetized rats on tone-shock association and found little conditioned emotional response (CER) when they were tested after waking; yet rats showed significant CER if they received epinephrine before and after the training session (Weinberger et al., 1984), and this memory lasted at least for a week (Gold et al., 1985). Such findings were supported by later studies showing trace conditioning in halothane-anesthetized mice (Pang et al., 1996) or discrimination learning in ketamine-treated rats (Edeline and Neuenschwander-El Massioui, 1988). However, some studies failed to find this effect on acquisition or extinction of nictitating membrane conditioning in rabbits treated by a sub-anesthetic dose of isoflurane (El-Zahaby et al., 1994) or on freezing conditioned to a tone in a context for rats under anesthesia of isoflurane or desflurane (Sonner et al., 2005).

Factors contributing to the discrepancy in the studies may include, among others, forms of learning or conditions of anesthesia and demand further elucidation (Samuel et al., 2018). The depth of anesthesia has been a major concern for observing such an effect or not (Lovibond and Shanks, 2002). The kind and dosage of anesthetics may affect this parameter that interacts with the action of epinephrine, since the latter is proposed to modulate memory by altering the arousal level (Gold and van Buskirk, 1975), which is closely related to the depth of anesthesia (Shin et al., 2004).

The way how memory was assessed may also contribute to the discrepancy. A previous study showed that humans could retrieve material learned under anesthesia via an implicit test as word stem completion but not via an explicit test as free recall (Deeprose et al., 2004). Animal studies with positive findings measured the memory indirectly by the fear-induced inhibition of a motivated behavior (water licking or bar pressing) (Weinberger et al., 1984; Gold et al., 1985; Edeline and Neuenschwander-El Massioui, 1988; Pang et al., 1996). The two studies with negative findings measured the memory directly by the defensive reaction (eye blink or freezing) elicited by the conditioned stimulus (CS) (El-Zahaby et al., 1994; Sonner et al., 2005). It is possible that the association acquired under anesthesia may express more readily in an indirect test (e.g. CER or saving test) than in a direct test.

In view of the above discussion, this study was designed to examine the effects of different anesthetics on different learning tasks. The effect of tone-shock conditioning under pentobarbital, an anesthetic acting on GABA_A_ and other receptors (Saunders and Ho, 1990), was assessed directly by the CS-induced freezing or indirectly by saving in re-learning. We also studied the light-shock association in a fear-potentiated startle task that assesses fear memory indirectly by the light-induced augmentation of acoustic startle (Brown et al., 1951). This task allows us to measure shock sensitivity during acquisition to monitor the anesthetic state. Dexmedetomidine is an anesthetic acting on α_2_ receptors of the locus coeruleus (Correa-Sales et al., 1992) and its action can be controlled in duration by an injection of antidote. We thus examined effects of this anesthetic on learning of the fear potentiated startle and the inhibitory avoidance tasks. The enhancing effect of epinephrine in awake rats, while is well documented in the inhibitory avoidance task (McGaugh, 1983), is less known for the other two tasks. As this could offer crucial information in searching for an epinephrine dose effective under anesthesia (Weinberger and Gold, 1995), we first examined in awake animals the epinephrine effect on the two tasks, and then explored possible shifts in the dose-response curve for epinephrine enhancement under anesthesia.

## Materials and Methods

### Subjects

Male Sprague-Dawley (SD) or Wistar rats used in this study were obtained from the National Animal Breeding Center or BioLASCO Company. Upon arrival at the lab in batches, they were housed in the university or department vivarium under a 12/12 light-dark cycle (lights on at 7:00 am) with 22∼27⁰C temperature and 50∼60% relative humidity. Rats were adapted to the lab for at least a month and had daily handling of 3 min for 5 days right before the experiment onset. The animal facility and experimental protocols followed regulation for animal studies of Taiwan Psychological Association and Council of Agriculture in Taiwan, ROC; and both were approved by Institutional Animal Care and Use Committees.

### Drug and Drug Administration

Groups of rats received an intra-peritoneal (i.p.) injection of sodium pentobarbital (50 mg/kg, Sigma, St. Louis, MO, USA) dissolved in 0.9% saline or a subcutaneous (s.c.) administration of dexmedetomidine (0.05 mg/kg, Dexdomitor®, Zoetis, USA) to initiate anesthesia about 0.5 h before training. Epinephrine hydrochloride (ampules of 1 mg/ml, China Chemical & Pharmaceutical, Taiwan) was diluted to an appropriate concentration by saline and injected s.c. either immediately after or 15 min before a training session. In animals undergoing a long training session, a priming dose (0.05 mg/kg) of dexmedetomidine was first injected and followed by continuous infusion of the drug at a rate of 1.0 ml/h (equivalent to a dose of 0.1 mg/kg/h) via an infusion pump to maintain stable anesthesia. After the training session, rats received a s.c. injection of atipamezole (0.8 mg/kg; Antisedan®, Zoetis, USA), the dexmedetomidine antidote, to terminate the anesthetic state.

### Behavioral Tasks

#### Conditioned freezing to tone and context

Conditioning to a tone and context was carried out in a Skinner box (30 × 24 × 24 cm, SG-501, MED Associates Inc., St. Albens, VT, USA) placed inside a wooden chamber (context A) or in the dark side of the inhibitory avoidance alley depicted in a latter section (context B). In each case, a loudspeaker was set up to emit auditory stimuli, and shocks were delivered through a shocker to the floor made up of 18 metal rods for context A or two metal plates with a slit in between for context B. The test box was either a plastic chamber (35 × 35 × 35 cm) with a trapezoid-shape interior created by a dark Plexiglas block with a floor of 24 bar rods (context C) or a wooden chamber (35 × 35 × 35 cm) with walls painted in white and a wire-mesh floor (context D). These test contexts were also equipped with lights for illumination and loudspeakers for delivery of auditory stimuli as in contexts for training.

Rats were placed into the training context, after 5 min of acclimation, a 5 kHz tone or white noise at 75 dB was presented as the CS for 30 s and in the last 2 s a 0.375/0.75 mA shock was delivered as the unconditioned stimulus (US), both were terminated together. Rats received 1 or 5 trials of CS-US pairing. One day after training, rats received two tests to assess conditioned freezing to the CS or context. Conditioned freezing was defined as lack of any body movement other than that of respiration. In the CS-specific test, rats were put into a test context different from that of training for 3-min without the tone (pre-CS) followed by 3-min presentation of the tone. The average of pre-CS freezing per minute was compared to the first minute of freezing in the presence of CS. Contextual conditioned freezing was tested 4 h later by counting the freezing behavior in the training context for a 6-min period. The rat’s behavior was recorded by video tape and the image was sampled subsequently every 4 s to code for the presence or absence of freezing behavior by a person blind to the treatment (Chang et al., 2008).

#### Conditioned fear potentiation of acoustic startle

This task was carried out in a set of startle apparatus (SR-LAB, San Diego Instruments, San Diego, CA, USA) in which a main module controlled four startle chambers. Each chamber was sound attenuated and air ventilated. The startle setup contained a cylindrical rat restrainer fixed on a platform underneath attached a sensor for vibration, which measured the maximal startle amplitude within 200 ms after the stimulus onset. Auditory stimuli of white noise or pure tone at various intensities were sent via a loud speaker located 30 cm above the restrainer. Electric shocks were delivered via the shock grid inserted into the restrainer connected to a shocker (TI 30, Colburn Instrument, Lehigh Valley, PA, USA). A light bulb was installed on the chamber wall. Presentation of all stimuli was controlled by programs set in the main module.

Rats underwent a matching process for the acoustic startle response. Each rat was placed into the apparatus with a 55-dB background noise. After acclimation for 5 min, 30 noise bursts (40 ms) of 95, 105 or 115 dB were delivered in a quasi-random order with 10 in each intensity and 30-s inter-burst intervals. The mean startle response for a rat was used to divide rats into groups with equivalent mean startle scores in each run of the experiment.

One day after the matching, rats received a fear conditioning procedure of light-shock pairing in an awake or anesthetic state. The rat was put into the restrainer with a shock grid. After 15-min adaptation to a 55-dB background noise, light-shock pairing was presented: The CS light was turned on for 3.5 s and the US shock appeared at the final 0.5 s with both terminating together. The inter-trial interval was 120 s. The total trial number would be specified in each experiment. The startle response to the shock was recorded as an index of shock sensitivity to monitor the anesthetic state.

The CS-induced potentiation of acoustic startle was tested 24 h after the training session in an awake state. The rat was placed into the apparatus and after 15 min of acclimation it received 70 trials. The first 10 trials were only noise bursts to stabilize the startle response. The remaining 60 trials were divided into 30 noise-alone trials and 30 light-noise trials. The former presented only noise bursts (either 100 dB or 95, 105 and 115 dB, specified in each experiment) and the latter presented first the light CS followed by the noise burst with the two terminating together. Both types of trials were presented with a 60-s inter-trial interval in a quasi-random order. If three sound intensities were used, each intensity was presented in one third of the trials.

The mean startle responses for each type of trials were obtained for all rats. The conditioned fear was represented by a percentage score indexing the amount of light-potentiation of acoustic startle for each rat. The raw percentage potentiation score was calculated as: [(mean startle score of the light-noise trials – mean startle score of the noise-alone trials)/mean startle score of the noise-alone trials] × 100%. As pretraining drug injections may sometimes alter the shock-sensitivity during the training session, the raw scores would be divided by a shock-sensitivity correction factor, which equals to the ratio of the shock-startle amplitude during training induced by a specific dose of epinephrine to that of saline. The correction yielded a rectified percentage potentiation score.

#### Inhibitory avoidance

The apparatus was a trough-shaped metal alley (100 cm long, 20 cm deep, 30 cm wide at the top and 6 cm wide at the floor) divided by a guillotine gate into lit and dark sides (40/60 cm), the latter was connected to a shocker (SMSCK-D Shocker, Kinder Scientific, Poway, CA, USA). The alley was installed in a longitudinal chamber. In the pre-exposure trial on Day 1, the rat was put into the lit side and allowed to explore the whole apparatus freely for 3 min. From Day 2 to Day 4, the rat was anesthetized by dexmedetomidine and 30 min later they were placed into the dark side of the apparatus to receive 10 shocks (1.2 mA/2 s at an inter-shock interval of 30 ± 5 s) with saline or epinephrine (0.1 mg/kg) injected 5 min before each of the three daily training sessions. The inhibitory avoidance response was tested on Day 5 in a waking state. The rat was put into the lit side of the alley facing against the gate, as it turned around the gate was opened to allow free entry into the dark side.

The latency (in seconds) from the gate opening to the rat’s stepping into the dark side with all four feet was recorded as a memory score. If a rat did not step in for 10 min, the test trial was terminated and a score of 600 s was assigned.

## Statistical Analysis

The freezing (experiment 1) and percentage potentiation scores (experiment 2 to 4) were subjected to parametric statistics, the overall differences among groups were detected by analyses of variance (ANOVAs). Comparisons between group pairs were accomplished by the LSD post-hoc analysis or a planned t test. As the distribution of inhibitory avoidance scores (experiment 5) was truncated at the 600 s ceiling, medians and inter-quartile ranges were used to represent the central and dispersion tendencies, respectively. Differences between groups were analyzed by the non-parametric Mann-Whitney U test.

## Results

### Experiment 1. Effects of epinephrine on conditioned freezing to tone and context

A total of 56 SD rats were divided into 4 groups and all received one tone-shock trial in context A, the shock was 0.375 mA/2 s. Rats received an injection of saline or epinephrine (0.01, 0.1 or 1.0 mg/kg) immediately after training. The results are shown in Table 1. The conditioned response to CS is indexed by the difference in freezing between the presence and absence of CS in context C. A two-way ANOVA revealed that the CS main effect was significant (*F*(1, 52) = 39.40, *p* < .01), indicating more freezing in the presence of CS for all groups. The main and interaction effects of the epinephrine variable were not significant (both *F*(3, 52) < 1), suggesting lack of any epinephrine enhancement. A one-way ANOVA of conditioned freezing in the context did not find a main effect of epinephrine (*F*(3, 52) < 1), suggesting that in an awake state epinephrine did not enhance conditioned freezing to the context.

**Table 1.** Conditioned freezing of rats trained in an awake or anesthetic state.

| Groups<br>(subjects) | Freezing (%; Mean ± SEM) | | |
| --- | --- | --- | --- |
|  | CS-absence | CS-presence | context |
| Awake State: 1-trial learning |  |  |  |
| Saline (24) | 9.9 ± 3.4 | 34.2 ± 5.9** | 22.7 ± 5.5 |
| 0.01 mg/kg (12) | 6.1 ± 3.0 | 33.3 ± 9.0** | 16.4 ± 3.5 |
| 0.1 mg/kg (9) | 4.7 ± 1.4 | 19.3 ± 8.9<sup>#</sup> | 24.2 ± 7.0 |
| 1.0 mg/kg (11) | 5.1 ± 2.1 | 23.6 ± 8.5* | 18.3 ± 6.4 |
| Pentobarbital Anesthesia: 1-trial / 5-trial learning |  |  |  |
| Saline (7) | 2.5 ± 1.2 / 8.6 ± 5.0 | 3.8 ± 2.0 / 4.8 ± 4.8 | 3.0 ± 0.9 / 4.0 ± 1.3 |
| 0.01 mg/kg (7) | 4.4 ± 2.1 / 3.8 ± 1.7 | 11.4 ± 6.3 / 15.2 ± 12.1 | 3.0 ± 1.4 / 7.1 ± 3.0 |
| 0.1 mg/kg (6) | 1.1 ± 0.8 / 5.2 ± 2.3 | 11.1 ± 11.1 / 3.3 ± 3.3 | 4.1 ± 1.5 / 2.6 ± 1.4 |
| 1.0 mg/kg (7) | 2.5 ± 1.4 / 8.6 ± 3.3 | 5.7 ± 3.1 / 7.6 ± 4.0 | 6.8 ± 3.6 / 1.7 ± 1.0 |

\*\* $p < .01$ , \* $p < .05$ , <sup>#</sup>.10 $< p < .05$ , different from the CS-absence condition

A group of 27 SD rats were used to test the epinephrine effect under anesthesia. Rats were first pre-exposed to context A for 2 min of free exploration. One day later, rats received a tone-shock (0.75 mA/2 s) pairing trial in context A under anesthesia of pentobarbital. Rats received an injection of epinephrine at 0.01, 0.1 or 1.0 mg/kg or saline immediately after training. Conditioned freezing to the tone and context was tested 1 day later in context C and A, respectively. Results are shown in Table 1. A two-way ANOVA showed that the difference in freezing before and after presentation of the CS approached statistical significance (*F*(1, 23) = 3.02, *p* = .10). Further, the various doses of epinephrine failed to enhance this weak learning as both the main and interaction effects of epinephrine were not significant (*F*(3, 23) < 1). As for the contextual conditioning, a one-way ANOVA indicated that epinephrine at the adopted doses failed to affect the conditioned freezing behavior in the 6-min test period (*F*(3, 23) < 1). These results showed that epinephrine did not affect the 1-trial learning of conditioned freezing under pentobarbital.

Eight days after the first training and testing episode, rats were trained by pre-exposure to context B for exploration and then received 5 trials of white noise-shock pairing under anesthesia with variable inter-trial intervals (mean = 60 s). These rats received the same post-training treatment as before. One day later, rats were tested for conditioned freezing to the CS and context as previously described in context D and B, respectively. Similar low freezing and lack of an epinephrine effect were found for this second learning episode as the data shown in Table 1. An ANOVA revealed no significant main or interaction effect on freezing to the CS (all *F*(3, 23) < 1) and no main effect on freezing to the context (*F*(3,23) = 1.65, *p* > .10). Under pentobarbital, 5 training trials still failed to yield significant conditioning and epinephrine effects.

One day after the second test session, all rats were placed in context A again in an awake state and received one relearning trial of tone-shock pairing (5 kHz/75 dB and 0.2 mA/0.5 s) without any further treatment. Conditioned freezing to the tone and the context was tested 1 day later, respectively, in context C and A. Results are shown in Figure 1, a two-way ANOVA of the test data after the re-learning trial revealed a significant main effect for presentation of the CS (*F*(1, 23) = 26.86, *p* < .01). Both the main effect and interaction effect involving epinephrine approached significance (*F*(3, 23) = 2.83 & 2.58, .05 < *p* < .08; respectively), yet post-hoc analyses revealed that the 0.01 or 0.1 mg/kg group showed more freezing in the CS presence than in its absence (*p* < .01); further under the CS presence, these two groups had higher freezing than the saline group (*p* < .05 & .01). Epinephrine also tended to enhance conditioned freezing to the context despite the difference fell short of statistical significance (*F*(3, 23) = 2.24, *p* = .11), yet post-hoc paired comparisons showed that freezing in the 0.1 mg/kg group was higher than that in the saline and 1.0 mg/kg groups (*p* < .05).

**Figure 1.**
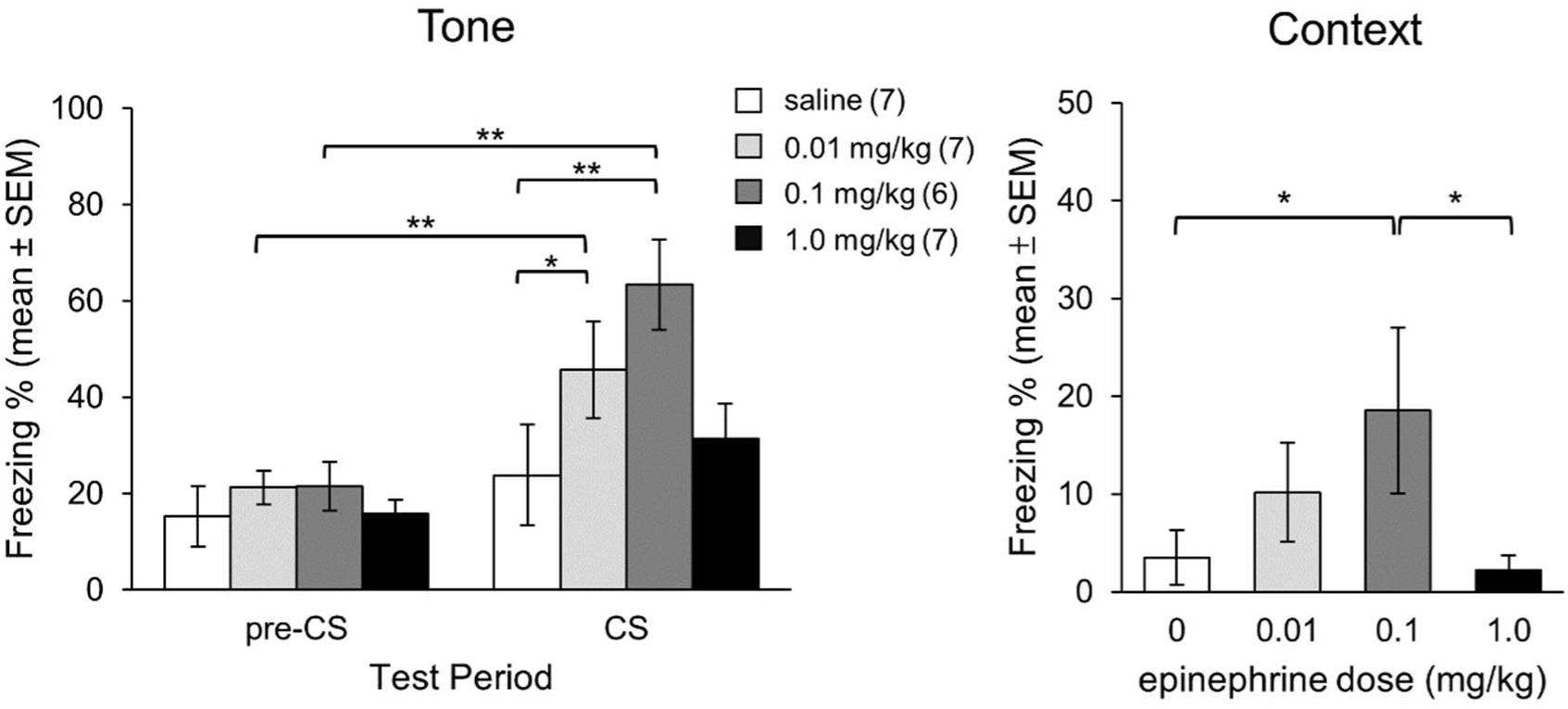
Rats trained under pentobarbital anethesia and treated with saline showed no conditioned freezing after retraining in an awake state. Yet epinephrine at 0.01 and 0.1 mg/kg given in the previous training episodes under anesthesia facilitated conditioned freezing to the tone in the retraining test and at 0.1 mg/kg facilitated conditioned freezing to the context in the retraining test. Number in the parethesis denotes n in each group. \**p* < .05, \*\**p* < .01.

### Experiment 2. Effects of epinephrine on one-trial training for conditioned fear potentiation of acoustic startle under awake and pentobarbital anesthesia

A group of 36 SD rats were trained in an awake state and another 36 were trained under pentobarbital anesthesia. They received one of the following treatments 15 min before the 1-trial light-shock paired training: saline or epinephrine in 0.01, 0.1 or 1.0 mg/kg. Fear memory was measured 1 day later in a waking state as described before. Results are shown in Figure 2. Figure 2A indicates that the shock startle in training appeared not to be the same for various groups but the difference was not statistically significant (*F*(3, 32) = 1.44, *p* > .20). For the raw percentage potentiation scores, the main effect of epinephrine was significant (*F*(3,32) = 3.46, *p* < .05), post-hoc group comparisons showed that the 0.01 or 0.1 mg/kg group had stronger potentiated startle than the saline group (*p* < .05), indicative of memory enhancement. If the raw scores were rectified by the shock-sensitivity correction factor as described in Materials and Methods, the results maintained that the 0.01 and 0.1 mg/kg groups had higher scores than the saline group based on the planned group comparisons (*t*(18) = 5.73 and 4.22, respectively, both *p* < .01), even though the main effect of epinephrine approached statistical significance only (*F*(3,32) = 2.50, *p* = .08). The 1.0 mg/kg group had large variability that rendered its difference from the saline group insignificant statistically.

**Figure 2.**
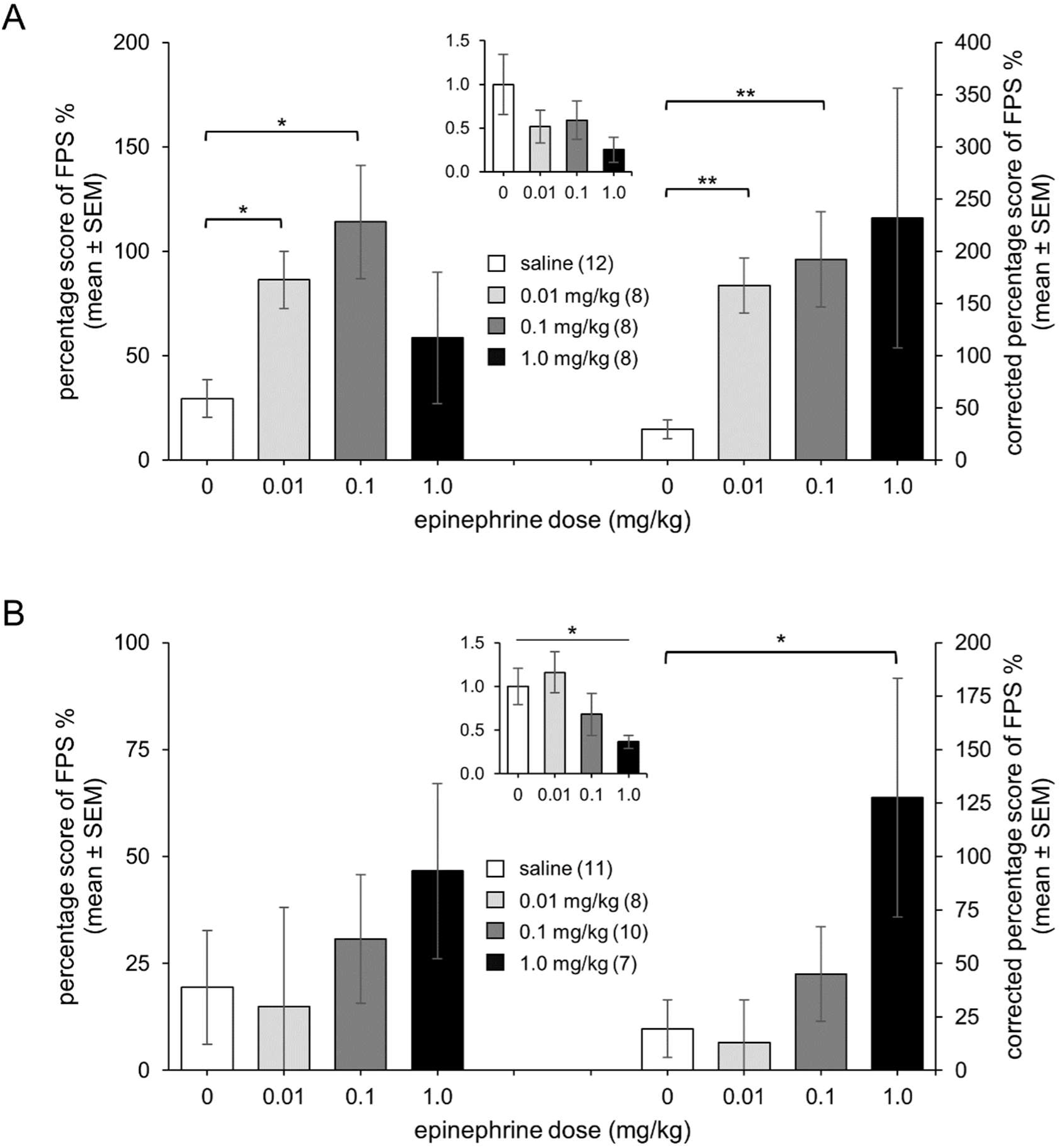
The effect of epinephrine on 1-trial learning of the fear potentiated startle (FPS) task. The left column shows the raw percentage scores of FPS and the right one shows the FPS scores rectified by a correction factor of the shock sensitity relative to the saline group during training (shown in the middle inset). (A) Rats trained in an awake state receving epinephrine at 0.01 or 0.1 mg/kg showed better fear memory than the saline controls for both the types of the FPS scores. (B) Rats receiving pentobartibal plus 1.0 mg/kg epinephrine showed a lower shock startle during training, but their rectified percentage scores of FPS were higher than those of the saline group. Number in the parethesis denotes n in each group. \**p* < .05, \*\**p* < .01.

Rats trained under anesthesia went through the same 1-trial light-shock pairing except for having 50 mg/kg pentobarbital 20 min before the epinephrine injection. The data are shown in Figure 2B. The shock startle during training under anesthesia seemed to be lower than the saline controls for rats receiving 1.0 mg/kg epinephrine but the difference only approached significance (*F*(3, 32) = 2.37, *p* = .09), a planned t test revealed a difference between the 1.0 mg/kg and saline groups (*t*(16) = 2.37, *p* < .05). When rats were tested in an awake state, the main effect of epinephrine was not significant (*F*(3,32) < 1). In view of a potential influence of epinephrine on the shock sensitivity during training, the raw percentage scores were rectified by a shock sensitivity factor of as previously described. A one-way ANOVA of the rectified scores yielded a significant epinephrine effect (*F*(3, 32) = 3.11, *p* < .05). Comparisons on the group pairs showed that the rectified percentage potentiation scores of the 1.0 mg/kg group was higher than those of the saline group (*p* < .05).

### Experiment 3. Effects of epinephrine on five-trial training for conditioned fear-potentiation of acoustic startle under awake and pentobarbital anesthesia

This experiment used a total of 123 SD rats to examine the epinephrine effect on conditioned-fear potentiation of startle with 5 light-shock paired trials under an awake or anesthetized state, it also examined the effect with 5 light-and-shock unpaired trials in anesthesia. The procedure and parameter were the same as experiment 2 except for rats receiving 5 training trials.

The first part of the experiment put 52 rats into four groups to receive an injection of saline or epinephrine at a dose of 0.01, 0.1 or 1.0 mg/kg 15 min before training in an awake state. Rats injected with 1.0 mg/kg epinephrine had the lower shock startle during training and a one-way ANOVA revealed a significant main effect (*F*(3, 48) = 3.50, *p* < .05). According to the post-hoc analyses, rats injected with 1.0 mg/kg epinephrine had lower shock-elicited startle than the 0.01 and 0.1 mg/kg group (*p* < .01). One day after training rats were tested for conditioned fear potentiation of startle and results are shown in Figure 3A. A one-way ANOVA revealed a significant main effect of epinephrine (*F*(3,48) = 4.64, *p* < .01), and post-hoc analysis indicated significant lower percentage scores of potentiation for the 1.0 mg/kg group (*p* < .05). In view of that the 1.0 mg/kg group showed lower shock sensitivity during training, we adjusted the raw percentage score by the correction factor as before. A one-way ANOVA on the rectified percentage scores of the fear potentiation of startle fell short of significance on the epinephrine main effect (*F*(3, 48) = 2.13, *p* > .10). However, a planned group comparison indicated that the difference between the 1.0 mg/kg and saline groups approached statistical significance (*t*(25) = 1.86, *p* = .08), indicative of an impairing trend.

**Figure 3.**
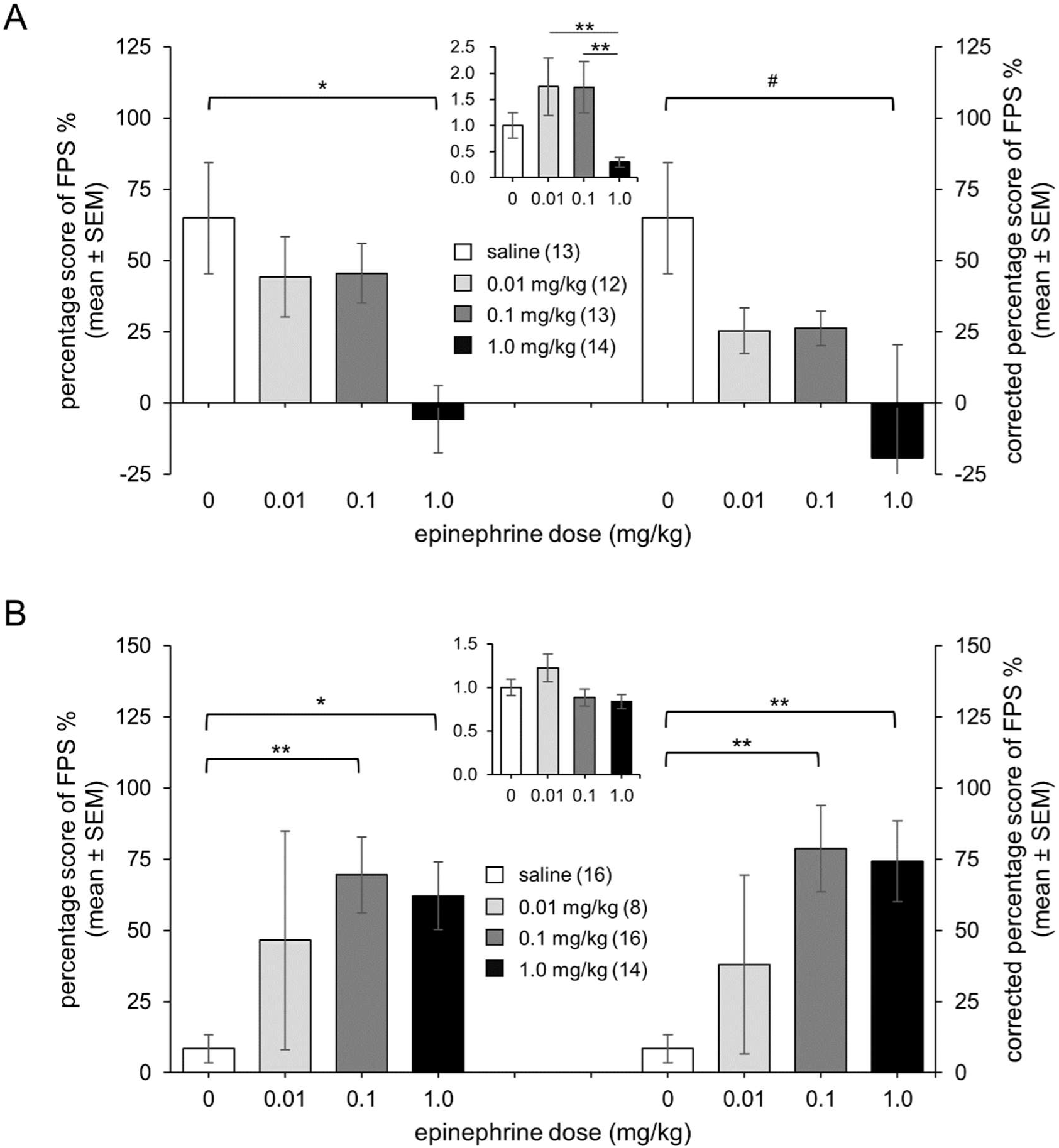
The effect of epinephrine on 5-trial learning of the fear potentiated startle (FPS) task. The left column shows the raw percentage score of FPS and the right one shows the FPS scores rectified by a correction factor of the shock sensitity relative to the saline group during training (shown in the middle inset). (A) Rats trained in an awake state showed lower shock startle during training and impaired FPS score in the group receving epinenphrine of 1.0 mg/kg. (B) Rats trained under pentobartibal anesthesia showed insignificant difference in shock startle during training. Epinephrine at 0.1 and 1.0 mg/kg facilitated learning under anesthesia for both raw and rectified percentage scores of FPS. Number in the parethesis denotes n in each group. \**p* < .05, \*\**p* < .01, ^#^.05 < *p* < .10.

The second part of this experiment trained 54 rats under pentobarbital anesthesia. The various groups injected with saline or epinephrine had similar shock-induced startle and a one-way ANOVA failed to detect a significant main effect of dose (*F*(3, 50) = 2.10, *p* > .10). Rats were assessed for conditioned fear-potentiation of startle in an awake state 24 h later and results are shown in Figure 3B. A one-way ANOVA revealed a significant main effect of epinephrine (*F*(3, 50) = 3.72, *p* < .05). Post-hoc analyses showed that the 0.1 and 1.0 mg/kg groups had higher percentage scores than the saline group (*p* < .01 & .05). Adjusting the raw percentage score by the correction factor of shock sensitivity as in the first part did not alter the pattern of results, a one-way ANOVA on the rectified percentage scores showed a significant effect among the various groups (*F*(3, 50) = 5.48, *p* < . 01) with the 0.1 and 1.0 mg/kg groups showing higher potentiation than the saline group (*p* < .01).

In the final part of this experiment, 17 anesthetized rats divided into two groups receiving saline or 0.1 mg/kg epinephrine were trained and tested as in other parts of this experiment except that the light and shock were presented in an explicit un-paired manner with an inter-stimulus interval of 60 s. The training data shown in Table 2 indicated that 0.1 mg/kg epinephrine did not influence the shock startle much under anesthesia (*F*(1, 15) = 1.49, *p* > .10). Five trials of unpaired training yielded very low percentage score of potentiated startle and epinephrine had negligible effect on it, one-way ANOVAs detected no significant difference between the two groups on either the raw or rectified percentage scores (*F*(1, 15) < 1 for both comparisons).

**Table 2.** Anesthetized rats trained with explicitly unpaired trials (Mean ± SEM)

| Groups (subjects) | Shock startle* | FPS % | rectified FPS % |
| --- | --- | --- | --- |
| saline (8) | 5.8 ± 1.0 (1.0 ± 0.2) | 16.7 ± 7.5 | 16.7 ± 7.5 |
| 0.1 mg/kg (9) | 7.2 ± 0.7 (1.2 ± 0.1) | 9.8 ± 8.8 | 7.8 ± 7.1 |

\*number in the parenthesis denotes the correction factor for shock sensitivity relative to the saline group.

### Experiment 4. Effect of epinephrine on conditioned fear potentiation of acoustic startle under dexmedetomidine anesthesia

This experiment used 48 Wistar rats to examine the effect of epinephrine on learning under dexmedetomidine. Rats received this anesthetic before and during the 20 light-shock training trials. A 2 × 2 factorial design was used to examine effects of epinephrine (saline or 0.1 mg/kg) and shock intensity (0.63 or 1.25 mA). The data are shown in Figure 4A. During training, light activated little startle and shock evoked a small but measurable response. The startle responses showed significant main effects of the shock intensity (*F*(1, 36) = 24.52, *p* < .01) and drug (*F*(1, 36) = 5.97, *p* < .05) during training, post-hoc analyses showed that 1.25 mA induced a stronger response than 0.63 mA (*p* < .01) and rats treated with epinephrine showed a smaller response than those treated with saline under strong shock (*p* < .05). An extra group trained by 0.63 mA foot shock and injected with 0.5 mg/kg epinephrine was included to examine whether a higher dose could yield a better effect on learning under dexmedetomidine anesthesia. The difference in the shock startle to 0.63 mA between this and the saline groups only approached statistical significance (*t*(13) = 1.86, *p* < .09).

**Figure 4.**
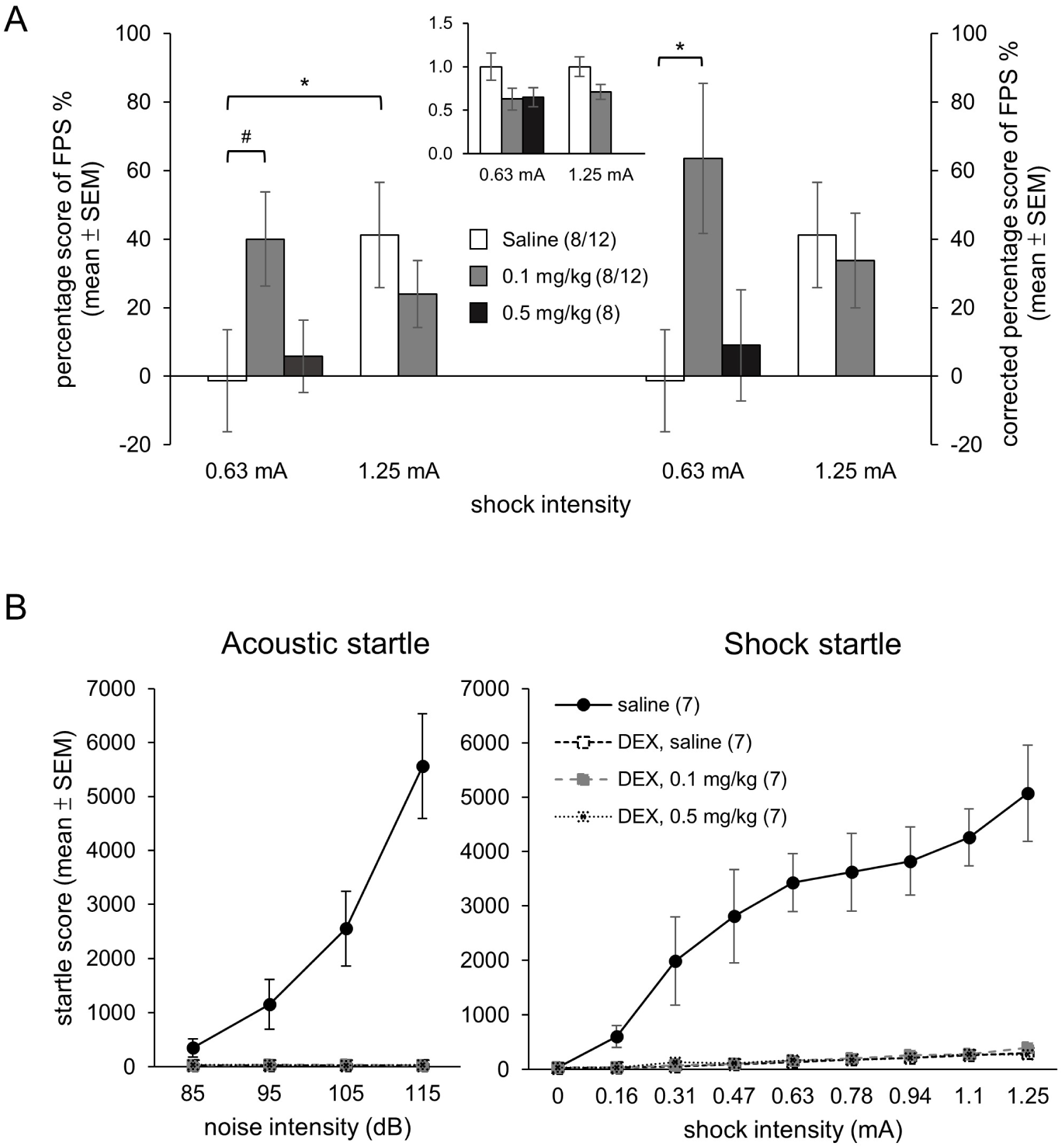
The effect of epinephrine on 20-trial learning of the fear potentiated startle (FPS) task under dexmedetomidine anethesia. (A) A shock intensity of 1.25 mA yielded better memory than that of 0.63 mA, but epinephrine at 0.1 mg/kg facilitated learning of rats trained with 0.63 mA. The left and right panels show, respectively, the raw FPS percentage scores and the rectified FPS percentage scores after correcting the shock sensitity relative to the saline group (shown in the middle inset). (B) Dexmedetomidine greatly reduced the amplitude of startle responses to either the auditory or shock stimuli, but epinephrine at no dose changed the state of dexmedetomidine anesthesia in this prolonged session as indicated by the startle to either the auditory or shock stimuli. Number in the parethesis denotes n for the 0.63 mA/1.25 mA group; \**p* < .05, ^#^.05 < *p* < .10.

Rats were tested in an awake state for conditioned fear potentiation of startle at the 95, 105, or 115 dB sound level. However, as the 115-dB stimulus often elicited a relatively high startle response in noise-alone trials, it left little room for potentiation in light-noise trials and thus yielded a low percentage score deviant greatly from those at the other two sound levels. Therefore, the analysis included only the data from 95 and 105 dB that had a mean of 100 dB, comparable to the test stimulus in the previous two experiments. The data shown in Figure 4A indicated that rats injected with saline and trained by 0.63 mA shock had little potentiation, but the saline-treated rats trained with 1.25 mA shock showed moderate potentiation. Pretraining administration of 0.1 mg/kg epinephrine enhanced the potentiation under the 0.63 mA shock but not under the 1.25 mA shock. A two-way ANOVA detected no significant main effects for the shock or epinephrine factor (both *F*(1, 36) < 1) but a significant interaction effect between the two (*F*(1, 36) = 4.42, *p* < .05). Post-hoc analyses showed that for rats injected with saline, the 1.25 mA group had higher percentage scores of potentiation than the 0.63 mA group (*p* < .05) and epinephrine tended to enhance the score in rats trained with the 0.63 mA shock as the difference between the epinephrine and saline groups approached significance (*p* = .06). Further analyses on the rectified percentage scores showed a significant interaction effect between the shock and epinephrine (*F*(1, 36) = 4.70, *p* < .05), paired comparisons indicated that rats treated with epinephrine of 0.1 mg/kg had higher rectified scores than the saline controls in the 0.63 mA condition (*p* < .05) but not in the 1.25 mA condition (*p* > .70). Further, epinephrine at a dose of 0.5 mg/kg did not have a facilitating effect on either the raw or rectified potentiation scores in the 0.63 mA condition (both *t*(13) < 1).

To study whether the epinephrine injection or repeated shock presentation may alter the anesthetic state of dexmedetomidine, additional 28 Wistar rats were divided into four groups to receive one of the following treatments: saline, dexmedetomidine plus saline, 0.1 or 0.5 mg/kg epinephrine given 15 min before presenting the stimulus. Three series of acoustic stimuli (85, 95, 105, 115 dB, twice for each intensity) and electric shocks (0, 0.16, 0.31, 0.47, 0,63, 0.78, 0.94, 1.10, 1.25 mA) were presented with 30 s inter-stimulus intervals after 15 min acclimation. The task period lasted for about 41 min. The stimuli of each series were arranged in a quasi-random order with the acoustic stimuli always preceding shocks. The data are shown in Figure 4B. The startle scores increased along with the stimulus intensity in the awake rats but not in the anesthetic ones; the difference among groups was significant in both acoustic and shock startle (*F*(3,24) = 22.83, *p* < .01; *F*(3,24) = 30.54, *p* < .01, respectively). The anesthetized rats did not respond much to the auditory stimuli, as their acoustic startle did not differ from that recorded in the background noise, in contrast, body movement to shocks was measurable. To analyze the difference in the conditions relevant to our training specifically, planned t tests were applied. The results showed that 1.25 mA shock did elicit stronger startle than 0.63 mA (*t*(6) = 4.22 & 5.71, both *p* < .01 for the dexmedetomidine groups receiving saline or 0.1 mg/kg epinephrine), but epinephrine did not significantly alter rats’ startle to the two shock levels in the series (*t*(13) = 0.91 and 1.55, respectively, for 0.63 and 1.25 mA, both *p* > .10). These results indicated that epinephrine did not significantly reduce the anesthetic state over a long session of sensory stimulation in different modalities and intensities.

### Experiment 5. Epinephrine enhanced inhibitory avoidance learning under dexmedetomidine anesthesia

Experiments 1 to 4 showed that with epinephrine facilitation rats acquired two classical conditioning tasks in anesthesia. Evidence has shown a pronounced effect of epinephrine on the inhibitory avoidance task (McGaugh, 1983), we thus examined if the effect was also present for learning this task under anesthesia. Based on the results of previous experiments, a total of 44 SD rats were trained on a two-stage inhibitory avoidance task with a 2 × 2 factorial design: Two groups of rats were pre-exposed to the apparatus on the first day, and two other groups were not; then one group in each condition received saline and the other one received 0.1 mg/kg epinephrine on the second day before having shocks in the dark side of the apparatus under anesthesia of dexmedetomidine. Figure 5 showed the results indicating that the two groups with no pre-exposure experience had low inhibitory avoidance scores, while the epinephrine-treated rats appeared to have higher scores than the saline controls, the difference only approached statistical significance (*U* = 31, .05 < *p* < .10). In contrast, for the two groups with the pre-exposure experience, the saline group showed poor inhibitory avoidance memory, and the epinephrine group showed better inhibitory avoidance memory than the saline group (*U* = 7.5, *p* < .001) and the epinephrine group witout pre-exposure (*U* = 29.5, *p* < .01). These data indicated that rats with a pre-exposure experience could acquire the aversive association between the apparatus and shock under anesthesia and express avoidance behavior in an awake state if they had been injected with epinephrine before training.

**Figure 5.**
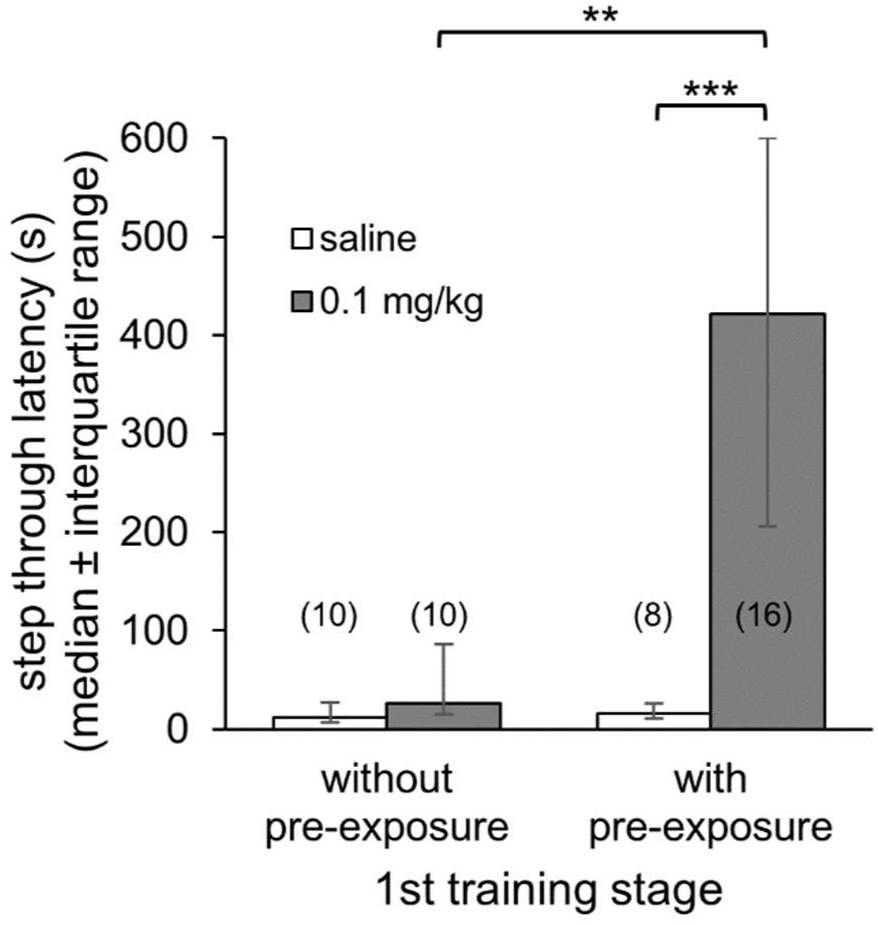
Epinephrine at 0.1 mg/kg injected before each of the three training sessions under dexmedetomidine anesthesia facilitated inhibitory avoidance learning for rats with the pre-exposure experience. Number in the parethesis denotes n in each group; \*\**p* < .01, \*\*\**p* < .001.

## Discussion

The present findings can be recapitulated as follows: In a fear conditioning task acquired in an awake or anesthetic state, epinephrine at various doses did not affect conditioned freezing to the CS or context, but it improved saving if rats trained under anesthesia were retrained in wakefulness. In a conditioned-fear potentiation of startle task, epinephrine at low to medium doses enhanced learning in awake rats for the 1-trial learning but at a high dose impaired the 5-trial learning. Rats had poor memory when trained under the pentobarbital anesthesia and epinephrine enhanced the 1-and 5-trial learning at different doses. Under dexmedetomidine anesthesia, training by a moderate foot shock yielded poor memory and epinephrine could improve it; training by a high foot shock yielded better memory but that was impervious to an epinephrine effect. Further, epinephrine enhanced inhibitory avoidance memory acquired in a two-stage training paradigm in which rats received shocks under dexmedetomidine at the second stage. These results suggest that epinephrine attenuated in a dose-dependent manner the suppressing effects of two anesthetics on memory of three learning tasks.

Consistent with the previous findings (Sonner et al., 2005), this study found no sign of conditioned freezing to the CS or context after learning under anesthesia and lack of any epinephrine effect in a direct test. Yet epinephrine enhanced saving in the retraining paradigm. These data support a proposal that an indirect test would be more sensitive to detect memory acquired under anesthesia and the superposed epinephrine effect. The conditioned-fear potentiation of startle also tests the memory indirectly by the CS potentiation of acoustic startle (Brown et al., 1951), analogous to the CER task that assesses fear memory by the CS inhibition of an ongoing behavior (Weinberger et al., 1984). The inhibitory avoidance response can be viewed as suppression of a pre-existing disposition of exploring the dark area by conditioned fear. A recent study also found that audio-visual association formed under anesthesia could modify appetitive or avoidance behavior in awake rats through second order conditioning (Zhang et al., 2020), indicating that memory formed in anesthesia could alter behavior tendencies in an awake animal. All these findings are consistent with those of human studies that implicit tests had better access to information acquired under anesthesia (Andrade, 1995; Deeprose et al., 2004; Deeprose and Andrade, 2006).

The present study monitored the anesthetic state during learning by the startle to shocks and found it to be much lower than that in the waking state and remained low consistently throughout the learning session under dexmedetomidine. Our informal observation also found that repeated foot shocks or epinephrine injections did not alter the state of pentobarbital anesthesia in a 90 min period (Chao, 2006). Thus, learning effects found in this study are unlikely due to instability of the anesthesia (Samuel et al., 2018). The epinephrine enhancement is not due to reduced anesthesia either, as epinephrine in the present study did not always elevate shock startle in anesthesia and we took such effects into account by correcting the raw scores with shock sensitivity, which did not alter the findings. Epinephrine injected before training in most of our experiments may affect the rats’ acquisition by altering their motivational state due to an aversive effect. Yet this is less likely to occur during anesthesia. Further, a recent study from this laboratory has ruled out this possibility by showing that epinephrine could enhance latent learning that involved no motivation (Liang and Chen, 2021).

Our results showed that epinephrine enhancement of learning under anesthesia depended on the doses and training conditions. In the 1-trial training on the fear potentiation of startle task, epinephrine enhanced memory at 0.01 and 0.1 mg/kg in a waking state but did so at 1.0 mg/kg under pentobarbital. In the 5-trial paradigm of same task learned by waking rats, epinephrine had no enhancing effect or even tended to impair memory at 1.0 mg/kg, but it enhanced learning at 0.1 and 1.0 mg/kg under pentobarbital. This shift of effective doses is explained by a view that epinephrine modulates memory by contributing to the general arousal level necessary for learning (Hebb, 1955; Konorski, 1967). It was proposed that epinephrine injected exogenously joins that released endogenously by the foot shock in contributing to the arousal level, which bears an inverted U function on memory consolidation (Gold, 2014). With the reduced brain arousal level by hypnotic or anesthetic drugs (Shaw et al., 2001; Kuo et al., 2009), the epinephrine enhancing dose would shift to a higher one for the 1-trial paradigm, and an impairing dose in the descending limb of the inverted-U function would become facilitating for the 5-trial paradigm. The shift may explain why some studies did not find an effect of epinephrine under anesthesia by testing only a limited range of doses based on findings in an awake state (El-Zahaby et al., 1994; Sonner et al., 2005).

The above interpretation may also account for our findings that under infusion of dexmedetomidine, 0.1 mg/kg epinephrine enhanced the multi-trial learning with 0.63 mA foot shocks but had no effect on that with 1.25 mA foot shocks, which replicated the previous ones that epinephrine at low doses enhanced memory trained with a low foot shock but had no effect with a high foot shock (Gold and van Buskirk, 1978). The better memory caused by 1.25 mA shocks than 0.63 mA shocks under anesthesia suggests that the sensory input evoked by multiple high shocks under anesthesia may be able to yield learning without supplementing epinephrine, just as found by the two studies adopting multi-trials of high shock training (Edeline and Neuenschwander-El Massioui, 1988; De Oliveira et al., 2020). Alternatively, it remains plausible that the release of endogenous epinephrine mediated by the spinal reflex under anesthesia may support such learning (Budgell et al., 1997). The endogenous release at high shocks may also render the injected epinephrine ineffective because the total amount of the hormone may be over the peak of the inverted-U curve, just as 0.5 mg/kg epinephrine failed to facilitate memory under the 0.63 mA foot-shock condition.

Our study also found that epinephrine facilitated an inhibitory avoidance learning task under dexmedetomidine in a two-phase multi-session training paradigm. Because epinephrine did not enhance learning under anesthesia by omitting the exploration at the first stage, this fear memory could be established by integrating a safe context memory formed during the exploration in an awake state with a shock memory under anesthesia. Our previous evidence has shown that this integrating process may involve the hippocampus to form a configural memory (Chang and Liang, 2017; Liang et al., 2019), which could support flexible application if rats are tested in situations different from the training (Chien et al., 2008). An involvement of the hippocampus in learning under anesthesia is implicated by hippocampal activity changes in trace conditioning of tone-shock association under urethane (De Oliveira et al., 2020) or the relationship between altered hippocampal Arc expression and learning under sevoflurane (Li et al., 2011).

It should be noted that epinephrine is not the only neurohumoral factor enabling acquisition of information under anesthesia. A recent study showed that peripheral or central administration of the cholecystokinin (CCK) analogs or stimulating the CCK terminals projecting from the entorhinal cortex to the auditory cortex enabled visual to auditory association under ketamine/xylazine anesthesia and altered evoked responses of the auditory cortex to visual stimuli (Zhang et al., 2020). As CCK is also implicated in emotional or arousal functions (Richter et al., 2014), these results are consistent with the notion that activating the arousal system could compensate for the learning deficit in anesthesia. In view of the CCK projection fibers from the amygdala to the bed nucleus of stria terminalis (Giardino and Pomrenze, 2021) and involvement of the two regions in emotion modulation of memory (Liang et al., 1983; Liu et al., 2009), it would be interesting to learn whether epinephrine and CCK may work in series or in parallel to enable learning under anesthesia.

In conclusion, our results provide clear evidence for fear learning in anesthesia with or without the assistance of epinephrine. The exact neural mechanism underlying how epinephrine may influence learning in anesthesia remains to be speculated. It was shown that epinephrine worked via the amygdala noradrenergic fibers to modulate memory formation elsewhere in the brain (Liang et al., 1986; McGaugh, 2004; Chen and Williams, 2012). This may also hold for learning under anesthesia as some recent studies implicated the noradrenergic fibers of the basolateral amygdala and their influence on the hippocampus in effects of anesthetics on memory (Li et al., 2011; Morena et al., 2021). Demonstration of learning under anesthesia allows on-line tracing of the neuronal plasticity associated with learning as shown by recent studies (De Oliveira et al., 2020; Zhang et al., 2020). Learning under dexmedetomidine, an anesthetic often used in animal research of functional brain imaging (Pawela et al., 2009; Chao et al., 2014; Sirmpilatze et al., 2019) is of interests in particular; as it offers an opportunity to map out plastic changes occurring in multiple brain regions during learning under anesthesia and thus allows correlating such widespread brain changes to the behavioral modification expressed by the animal in an awake state. This is what we pursued in the study to follow.

## Acknowledgment

Experiments in the present study was supported by Ministry of Science and Technology in grants 95-2413-H002-028-MY3, 101-2410-H002-MY3, 105-2410-H002-051 to KCL and 105-2410-H-006-017, 106-2410-H-006-037, 107-2410-H-006-054 to DYC. The authors would like to thank the students involved in part of the experiments of this study.

## References

Andrade, J. (1995). Learning during anesthesia: A review. Br J Psychol, 86, 479–506. DOI: 10.1111/j.2044-8295.1995.tb02566.x

Brown, J. S., Kalish, H. I., & Farber, L. E. (1951). Conditioned fear as revealed by magnitude of startle response to an auditory stimulus. J Exp Psychol, 41, 317–328. DOI: 10.1037/h0060166

Budgell, B., Sato, A., Suzuki, A., & Uchida, S. (1997). Responses of adrenal function to stimulation of lumbar and thoracic interspinous tissues in the rat. Neurosci Res, 28, 33–40. DOI: 10.1016/s0168-0102(97)01173-5

Chang, S. D., & Liang, K. C. (2017). The hippocampus integrates context and shock into a configural memory in contextual fear conditioning. Hippocampus, 27, 145–155. DOI: 10.1002/hipo.22679

Chang, S. D., Chen, D. Y., & Liang, K. C. (2008). Infusion of lidocaine into the dorsal hippocampus before or after the shock training phase impaired conditioned freezing in a two-phase training task of contextual fear conditioning. Neurobiol Learn Mem, 89, 95–105. DOI: 10.1016/j.nlm.2007.07.012

Chao, S. T. (2006). Epinephrine modulation of fear conditioning under awake and anesthetic states. Master thesis for Department of Psychology, National Taiwan University.

Chao, T. H. H., Chen, J. H., & Yen, C. T. (2014). Repeated BOLD-fMRI imaging of deep brain stimulation responses in rats. PLoS ONE, e97305. DOI: 10.1371/journal.pone.0097305

Chen, C. C., & Williams, C. L. (2012). Interactions between epinephrine, ascending vagal fibers and central noradrenergic systems in modulating memory for emotionally arousing events. Front Behav Neurosci, 6, 35. DOI: 10.3389/fnbeh.2012.00035

Chien, W. L., Liang, K. C., & Fu, W. M. (2008). Enhancement of active shuttle avoidance response by the NO-cGMP-PKG activator YC-1. Eur J Pharmacol, 590, 233–240. DOI: 10.1016/j.ejphar.2008.06.040

Correa-Sales, C., Rabin, B. C., & Maze, M. (1992). A hypnotic response to dexmedetomidine, an α2 agonist, is mediated in the locus coeruleus in rats. Anesthesiology, 76, 948–952. DOI: 10.1097/00000542-199206000-00013

De Oliveira, E. F., Dickson, C. T., & Reyes, M. B. (2020). Hippocampal and lateral entorhinal cortex physiological activity during trace conditioning under urethane anesthesia. J Neurophysiol, 124, 781–789. DOI: 10.1152/jn.00293.2020

Deeprose, C., & Andrade, J. (2006). Is priming during anesthesia unconscious? Conscious Cogn, 15, 1–23. DOI: 10.1016/j.concog.2005.05.003

Deeprose, C., Andrade, J., Varma, S., & Edwards, N. (2004). Unconscious learning during surgery with propofol anaesthesia. Br J Anaesth, 92, 171–177. DOI: 10.1093/bja/aeh054

El-Zahaby, H. M., Ghoneim, M. M., Johnson, G. M., & Gormezano, B. A. (1994). Effects of subanesthetic concentrations of isoflurane and their interaction with epinephrine on acquisition and retention of the rabbit nictitating membrane response. Anesthesiology, 81, 229–237. DOI: 10.1097/00000542-199407000-00029

Edeline, J. M., & Neuenschwander-el Massioui, N. (1988). Retention of CS–US association learned under ketamine anesthesia. Brain Res, 457, 274–280. DOI: 10.1016/0006-8993(88)90696-8

Giardino, W. J. & Pomrenze, M. B. (2021). Extended amygdala neuropeptide circuitry of emotional arousal: Waking up on the wrong side of the bed nucleus of stria terminalis. Front Behav Neurosci, 15, 613025. DOI: 10.3389/fnbeh.2021.613025

Gidron, Y., Barak, T., Henik, A., Gurman, G., & Stiener, O. (2002). Implicit learning of emotional information under anesthesia. Neuroreport, 13, 139–142. DOI: 10.1097/00001756-200201210-00032

Gold, P. E. (2014). Regulation of memory—From the adrenal medulla to liver to astrocytes to neurons. Brain Res Bull, 105, 25–35. DOI: 10.1016/j.brainresbull.2013.12.012

Gold, P. E., & van Buskirk, R. (1975). Facilitation of time-dependent memory processes with posttrial epinephrine injections. Behav Biol, 13, 145–153. DOI: 10.1016/s0091-6773(75)91784-8

Gold, P. E., & van Buskirk, R. (1978). Posttraining brain norepinephrine concentrations: correlation with retention performance of avoidance training and with peripheral epinephrine modulation of memory processing. Behav Biol, 23, 509–520. DOI: 10.1016/s0091-6773(78)91614-0

Gold, P. E., Weinberger, N. M., & Sternberg, D. B. (1985). Epinephrine-induced learning under anesthesia: Retention performance at several training-testing intervals. Behav Neurosci, 99, 1019–1022. DOI: 10.1037//0735-7044.99.5.1019

Hebb, D. O. (1955). Drives and the C.N.S. (conceptual nervous system). Psychol Rev, 62, 243–254. DOI: 10.1037/h0041823

Inostroza, M., & Born, J. (2013). Sleep for preserving and transforming episodic memory. Annu Rev Neurosci, 36, 79–102. DOI: 10.1146/annurev-neuro-062012-170429

Kim, E. H., Woody, C. D., & Berthier, N. E. (1983). Rapid acquisition of conditioned eye blink response in cats following pairing of an auditory CS with glabella tap US and hypothalamic stimulation. J Neurophysiol, 49, 767–779. DOI: 10.1152/jn.1983.49.3.767

Klinzing, J. G., Niethard, N., & Born J. (2019). Mechanisms of systems memory consolidation during sleep. Nat Neurosci, 22, 1598–1610. DOI: 10.1038/s41593-019-0467-3

Konorski, J. (1967). Integrative activity of the brain: An interdisciplinary approach. Chicago, IL: University of Chicago Press.

Kuo, C. C., Chiou, R. J., Liang, K. C., & Yen, C. T. (2009). Differential involvement of anterior cingulate and primary sensorimotor cortices in sensory and affective functions of pain. J Neurophysiol, 101, 1201–1210. DOI: 10.1152/jn.90347.2008

Lennartz, R. C., & Weinberger, N. M. (1992). Analysis of response systems in Pavlovian conditioning reveals rapidly versus slowly acquired conditioned responses: Support for two factors, implications for behavior and neurobiology. Psychobiology, 20, 93–119. DOI: 10.3758/BF03327169

Li, Q., Liu, X. S., Zeng, Q. W., Xue, Q. S., Cao, X. H., Liu, J., Ren, Y., & Yu, B. W. (2011). Post-training intra-basolateral amygdala infusions of norepinephrine block sevoflurane-induced impairment of memory consolidation and activity-regulated cytoskeletal protein expression inhibition in rat hippocampus. Neurobiol Learn Mem, 96, 492–497. DOI: 10.1016/j.nlm.2011.08.002

Liang, K. C. & Chen, D. Y. (2021). Epinephrine modulates memory of latent learning in an inhibitory avoidance task. Neurobiol Learn Mem, 182, 107447. DOI: 10.1016/j.nlm.2021.107447

Liang, K.C., Chen, D.Y., & Chang, S.D. (2019). A tale of two engrams: Roles of the hippocampus and amygdala in contextual fear conditioning. Chin J Psychol, 61, 321–340. DOI: 10.6129/CJP.201912_61(4).0004

Liang, K. C., Juler, R. G., & McGaugh, J. L. (1986). Modulating effects of posttraining epinephrine on memory: Involvement of the amygdala noradrenergic system. Brain Res, 368, 125–133. DOI: 10.1016/0006-8993(86)91049-8

Liang, K. C., Messing, R. B., & McGaugh, J. L. (1983). Naloxone attenuates amnesia caused by amygdaloid stimulation: The involvement of a central opioid system. Brain Res, 271, 41–49. DOI: 10.1016/0006-8993(83)91363-x

Liu, T. L., Chen, D. Y., & Liang, K. C. (2009). Posttraining infusion of glutamate into the bed nucleus of the stria terminalis enhanced inhibitory avoidance memory: an effect involving norepinephrine. Neurobiol Learn Mem, 91, 456–465. DOI: 10.1016/j.nlm.2009.01.003

Lovibond, P. F., & Shanks, D. R. (2002). The role of awareness in Pavlovian conditioning: Empirical evidence and theoretical implications. J Exp Psychol Anim Behav Process, 28, 3–26. DOI: 10.1037/0097-7403.28.1.3

McGaugh, J. L. (1983). Hormonal influences on memory. Annu Rev Psychol, 34, 297–323. DOI: 10.1146/annurev.ps.34.020183.001501

McGaugh, J. L. (2004). The amygdala modulates the consolidation of memories of emotionally arousing experiences. Annu Rev Neurosci, 27, 1–28. DOI: 10.1146/annurev.neuro.27.070203.144157

Merikle, P. M., & Daneman, M. (1996). Memory for unconsciously perceived events: Evidence from anesthetized patients. Conscious Cogn, 5, 525–541. DOI: 10.1006/ccog.1996.0031

Morena, M., Colucci, P., Mancini, G. F., De Castro, V., Peloso, A., Schelling, G., & Campolongo, P. (2021). Ketamine anesthesia enhances fear memory consolidation via noradrenergic activation in the basolateral amygdala. Neurobiol Learn Mem, 178, 107362. DOI: 10.1016/j.nlm.2020.107362

Nicol, A. U., Sanchez-Andrade, G., Collado, P., Segonds-Pichon, A., & Kendrick, K. M. (2014). Olfactory bulb encoding during learning under anesthesia. Front Behav Neurosci, 8, 193. DOI: 10.3389/fnbeh.2014.00193

Pang, R., Turndorf, H., & Quartermain, D. (1996). Pavlovian fear conditioning in mice anesthetized with halothane. Physiol Behav, 59, 873–875. DOI: 10.1016/0031-9384(95)02137-x

Pavlov, I. P. (1927). Conditioned reflexes: an investigation of the physiological activity of the cerebral cortex. Oxford, England: Oxford University Press.

Pawela, C. P., Biswal, B. B., Hudetz, A. G, Schulte, M. L., Li. R., Jones, S. R., Chao, Y. R., Matloub, H. S., et al. (2009). A protocol for use of medetomidine anesthesia in rats for extended studies using task-induced BOLD contrast and resting state functional connectivity. Neuroimage, 46, 1137–1147. DOI: 10.1016/j.neuroimage.2009.03.004

Reasor, J. D., & Poe, G. R. (2008). Learning and memory during sleep and anesthesia. Int Anesthesiol Clin, 46, 105–129. DOI: 10.1097/AIA.0b013e318181e513

Richter, C., Woods, I. G., & Schier, A. F. (2014). Neuropeptidergic control of sleep and wakefulness. Annu Rev Neurosci, 37, 503–531. DOI: 10.1146/annurev-neuro-062111-150447

Samuel, N., Taub, A. H., Paz, R., & Raz, A. (2018). Implicit aversive memory under anesthesia in animal models: A narrative review. Br J Anaesth, 121, 219–232. DOI: 10.1016/j.bja.2018.05.046

Saunders, P. A., & Ho, I. K. (1990). Barbiturates and the GABA_A_ receptor complex. Prog Drug Res, 34, 261–286. DOI: 10.1007/978-3-0348-7128-0_7

Shaw, F. Z., Chen, R. F., & Yen, C. T. (2001). Dynamic changes of touch- and laser heat-evoked field potentials of primary somatosensory cortex in awake and pentobarbital-anesthetized rats. Brain Res, 911, 105–115. DOI: 10.1016/s0006-8993(01)02686-5

Sirmpilatze, N., Baudewig, J., & Boretius, S. (2019). Temporal stability of fMRI in medetomidine-anesthetized rats. Sci Rep, 9, 16673. DOI: 10.1038/s41598-019-53144-y

Shin, H. W., Ban, Y. J., Lee, H. W., Lim, H. J., Yoon, S. M., & Chang, S. H. (2004). Arousal with iv epinephrine depends on the depth of anesthesia. Can J Anaesth, 51, 880–885. DOI: 10.1007/BF03018884

Sonner, J. M., Xing, Y., Zhang, Y., Maurer, A., Fanselow, M. S., Dutton, R. C. & Eger II, E. I. (2005). Administration of epinephrine does not increase learning of fear to tone in rats anesthetized with isoflurane or desflurane. Anesth Analg, 100, 1333–1337. DOI: 10.1213/01.ANE.0000148619.77117.C7

Weinberger, N. M., Gold, P. E., & Sternberg, D. B. (1984). Epinephrine enables Pavlovian fear conditioning under anesthesia. Science, 223, 605–607. DOI: 10.1126/science.6695173

Weinberger, N. M., & Gold, P. E. (1995). What is a “replication”? Epinephrine facilitation of learning under anesthesia. Anesthesiology, 82, 308–310. DOI: 10.1097/00000542-199501000-00040

Zhang, Z., Zheng, X., Sun, W., Peng, Y., Guo, Y., Lu, D., Zheng, Y., Li, X. et al. (2020). Visuoauditory associative memory established with cholecystokinin under anesthesia is retrieved in behavioral contexts. J Neurosci, 40, 2025–2037. DOI: 10.1523/JNEUROSCI.1673-19.2019

